# The FW2.2/CNR protein regulates cell-to-cell communication in tomato by modulating callose deposition at plasmodesmata

**DOI:** 10.1101/2023.09.27.559775

**Authors:** Arthur Beauchet, Norbert Bollier, Magali Grison, Valérie Rofidal, Frédéric Gévaudant, Emmanuelle Bayer, Nathalie Gonzalez, Christian Chevalier

## Abstract

The *FW2.2* gene is the founding member of the *CELL NUMBER REGULATOR* (*CNR*) gene family. More than 20 years ago, *FW2.2* was the first cloned gene underlying a Quantitative Trait Locus (QTL) governing fruit size/weight in tomato. However, despite this discovery, the molecular mechanisms by which FW2.2 acts as a negative regulator of cell divisions during fruit growth remain undeciphered. In the present study, we confirm that FW2.2 is a transmembrane spanning protein, whose both N- and C-terminal ends are facing the apoplast. We unexpectedly found that FW2.2 is located at plasmodesmata (PD). FW2.2 participates in the spatiotemporal regulation of callose deposition at PD via an interaction with Callose Synthases, which suggests a regulatory role in cell-to-cell communication by modulating PD transport capacity and trafficking of signaling molecules during fruit development.

## INTRODUCTION

The tight coordination of developmental processes such as cell division, cell expansion and cell differentiation, is pivotal for proper plant growth at the whole organismal, organ and tissue level. Unravelling the genes that contribute to impact plant yield and biomass, and improve agronomical quality traits, is thus a major goal of plant biology and agronomy. In the particular case of tomato fruit size determination, nearly 30 Quantitative Trait Loci (QTL) governing fruit size/weight have been identified (Grandillo et al., 1999; Lippman and Tanksley, 2001; van der Knaap and Tanksley, 2003). However, the molecular basis governing these QTLs remains mostly undeciphered, and only three major genes underlying such QTLs in tomato have been identified and cloned so far (Frary et al., 2000; Chakrabarti et al., 2013; Mu et al., 2017).

FW2.2 was the first cloned gene underlying a QTL related to fruit size in tomato (Alpert et al., 1995; Frary et al., 2000). The encoded protein FW2.2 was defined as a major negative regulator of cell divisions in young developing fruit, and thus impacting fruit size (Frary et al., 2000; Cong et al., 2002; Liu et al., 2003; Nesbitt and Tanksley, 2001; Baldet et al., 2006). FW2.2 was the founding member of the CELL NUMBER REGULATOR/FW2.2-Like (CNR/FWL) protein family (Guo et al., 2010), whose function in organ size control seems to be conserved in both monocotyledon and dicotyledon plants (for a review, see Beauchet et al., 2021). Members of this protein family possess a conserved PLAC8 (Placenta-specific gene 8 protein) domain (Galaviz-Hernandez et al., 2003), which is composed of one or two hydrophobic segments, predicted to form transmembrane (TM) helices (Song et al., 2004). The hydrophobic segments are characterized by the presence of conserved Cys-rich motifs of the type CLXXXXCPC or CCXXXXCPC, separated by a variable region and located at the N-terminal part of a first TM domain (Beauchet et al., 2021). A localization at the plasma membrane (PM) was indeed demonstrated for the tomato FW2.2 protein (Cong and Tanksley, 2006), as well as for CNR/FWL homologous proteins in various fruit species such as eggplant, pepper, Physalis, avocado, cherry (Dahan et al., 2010; De Franceschi et al., 2013; Doganlar et al., 2002; Li and He, 2015), but also in Arabidopsis, cereal and leguminous species (Libault et al., 2010; Guo et al., 2010; Song et al., 2010; Xu et al., 2013). In soybean, the CNR/FWL protein GmFWL1 was shown to display a punctate localization in plasma membrane nanodomains, which supported its ability to interact with membrane nanodomain-associated proteins such as flotillins, prohibitins, remorins, proton- and vacuolar-ATPases, receptor kinases, leucine-rich repeat proteins (Qiao et al., 2017).

Despite the seemingly conserved roles in cell division and organ size control (Beauchet et al., 2021), the precise physiological and biochemical function of FW2.2 or its CNR/FWL homologues remains unknown so far. The conceptual question in studying the functional role of FW2.2 and CNR/FWL is thus how to conciliate a localization at the plasma membrane and nanodomains with a spatial and temporal control of cell divisions in order to regulate plant organ growth.

In plants, important biological functions are associated to membrane nanodomains. Plasmodesmata (PD) belong to such PM nanodomains. PD are cell wall- and membrane-spanning channels, which provide direct cytosolic continuity to mediate symplastic communication between cells (Maule et al., 2011; Petit et al., 2020). PD control cell-to-cell movements of different mobile signalling molecules (Van Norman et al., 2011; Gallagher et al., 2014), and thus regulate the connection between cells ensuring both local and systemic responses to biotic and abiotic stresses, the exchange of nutrients and organs, regulating symbiotic interactions and supporting the coordination of developmental processes (Gaudioso-Pedraza et al., 2018; Grison et al., 2019; Han et al., 2014a; O’Lexy et al., 2018; Yan et al., 2019). Hormones, metabolites, non-cell autonomous proteins, including transcription factors (TFs), and small RNAs represent such mobile signalling molecules, trafficking from cell-to-cell via PD. The symplastic communication via PD is finely tuned by developmental or environmental factors, which exert a control on the size exclusion limit of PD. Among these factors, the deposition of callose, a (1,3)-β-glucan polymer, regulated by the antagonistic action of callose synthases and β-glucanases, is a major process that constricts the PD channel, and thus decreases the aperture of PD (Amsbury et al., 2018). Consequently, the balance between callose deposition and degradation at the neck region of PD plays a major role in the regulation of cell-to-cell communication.

In an effort to unravel the cellular and molecular mechanisms sustaining the mode of action of FW2.2 in tomato, we re-investigated its subcellular localization *in planta*. We unexpectedly found that FW2.2 protein not only associates with bulk PM but also clusters at PD in the different tissues we examined. We further show that FW2.2 modulates the functionality of PD by modifying callose levels. FW2.2-induced regulation of callose most likely occurs through direct interaction with PD-associated Callose Synthases. Our data shed light on an unforeseen function of FW2.2 in modulating cell-to-cell communication in tomato.

## RESULTS

### FW2.2 localizes at the plasma membrane with the N- and C-terminal parts facing the apoplast

The first and only demonstration that FW2.2 addresses the PM was provided by transient expression analysis using onion epidermal cells and tomato young leaf cells (Cong and Tanksley, 2006). This PM localization is conferred by the two transmembrane domains (TMD) contained in the PLAC8 domain, but the exact topology of the FW2.2 protein at PM is still uncharacterized.

First, we confirmed the PM localization of FW2.2, using transient expression in *Nicotiana benthamiana* leaves (Xie et al., 2017). FW2.2 fused to GFP either at its C-terminus of N-terminus was indeed addressed to the PM (**Figure 1A**). To investigate the mode of action of FW2.2 at PM, we then study the topology of FW2.2 by using a Bi-molecular Fluorescent Complementation (BiFC) approach that had been validated for PM-located proteins (Thomas et al., 2008). The FW2.2 protein was fused at its N- or C-terminus to the truncated version of GFP, namely GFP11, which contains the last and eleventh β-sheet. The GFP11-FW2.2 or FW2.2-GFP11 construct was then co-expressed with the cytosolic truncated version of the GFP, namely GFP1-10 containing the first ten β-sheets. Alternatively, the GFP11-FW2.2 or FW2.2-GFP11 construct was co-expressed with a secreted apoplastic version of GFP1-10, namely SP-GFP1-10 (SP for Signal Peptide of the Arabidopsis PR1 protein; At2g14610). As a positive control for a cytosolic interaction, we fused the GFP11 to the C-terminal part of the PM located protein Lti6b (Low-temperature induced 6b protein; At3g05890) that faces the cytosol (Martiniere et al., 2012), and co-infiltrated this construct with the GFP1-10. The Lti6b-GFP11 construct was thus expected to be unable to interact with the apoplastic SP-GFP1-10.

**Figure 1.**
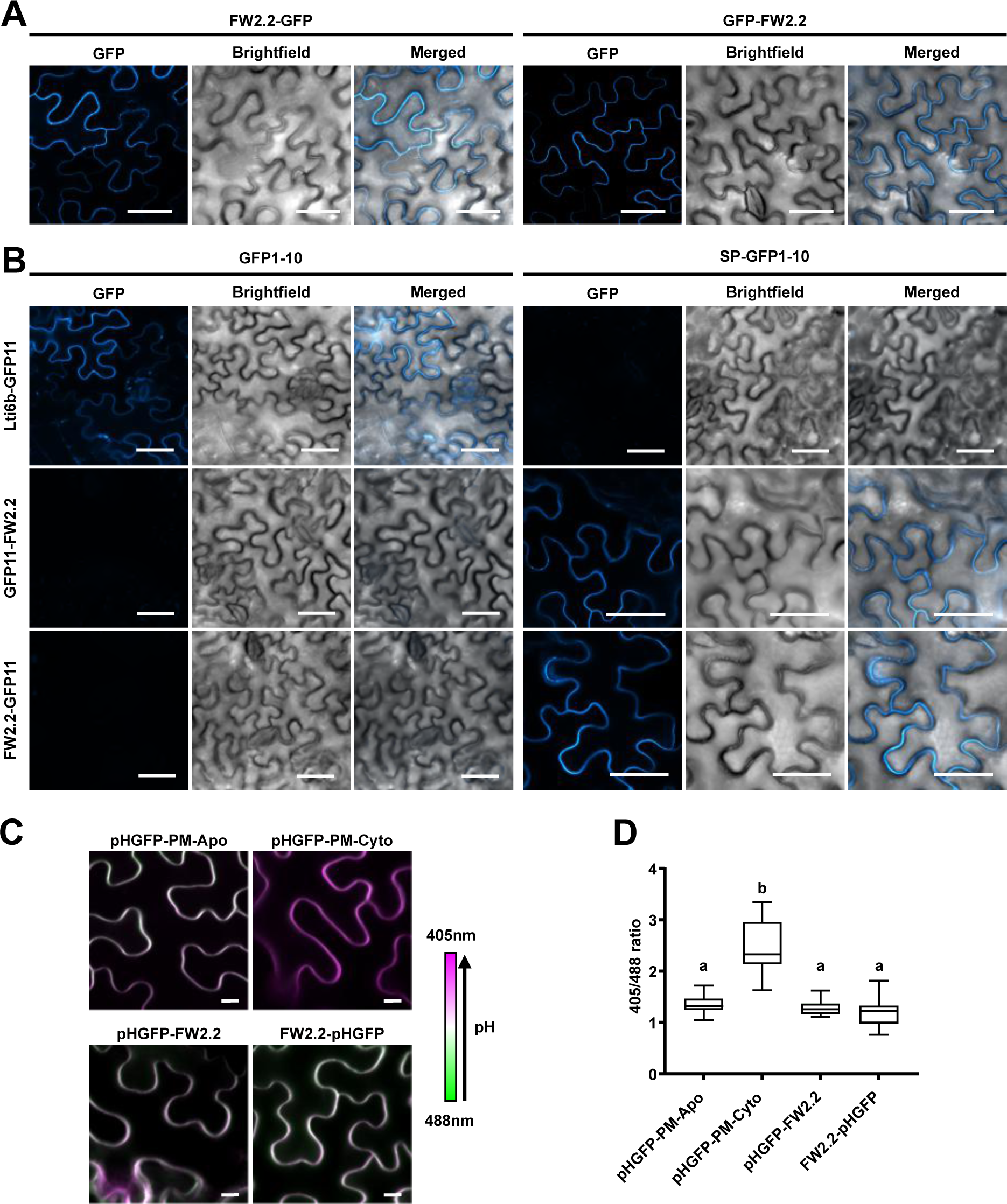
Topological analysis of FW2.2 at the plasma membrane. **(A)** Subcellular localization of FW2.2 fused to GFP in *N. benthamiana* leaf epidermal cells. **(B)** BiFC assays deciphering the topology of FW2.2 at the plasma membrane. Transient expressions of FW2.2 or Lti6b fused to GFP11 and with a cytosolic GFP (GFP1-10) or a apoplastic GFP (SP-GFP1-10) were performed in *N. benthamiana* leaves, followed by observation using confocal microscopy Scale bar = 50 µm. **(C)** Confocal imaging of pHGFP-PM-Apo, pHGFP-PM-Cyto and pHGFP fused to FW2.2 at the N- and C-terminus in *N. benthamiana* leaf epidermal cells. The four images were taken using the same confocal settings. Scale bar = 10µm. **(D)** 405/488 intensity ratio at plasma membrane. n>15. ANOVA followed by Tukey’s test; P < 0.05 between a and b groups.

A strong GFP signal was observed when the Lti6b-GFP11 was co-expressed with the cytosolic GFP1-10, and no signal was observed when co-expressed with the apoplastic SP-GFP1-10, which validated the BiFC approach (**Figure 1B**). The co-expression of FW2.2 fused to GFP11 at both its C- and N-terminus with the cytosolic GFP1-10, did not result in any visible fluorescence signal. On the contrary, the co-expression of FW2.2 fused to GFP11 with the apoplastic SP-GFP1-10 resulted in a strong GFP signal at the PM (**Figure 1B**). Therefore, we confirmed that FW2.2 is associated to PM as previously reported (Cong and Tanksley, 2006), but we provided evidence that the two TMDs within FW2.2 drives a protein topology where the N- and C-terminus are facing the apoplast.

To confirm this topology, we performed a second transient expression assays, using a system of apoplastic and cytoplasmic pH sensors described by Martinière et al. (2018). This system takes advantage of the pH-sensitive ratiometric behavior of the protein pHluorin (pHGFP), whose emitted fluorescence differs according to its location in the cytosol or the apoplast, depending on their respective pH value of ∼7.5 or ∼6.0. Following agro-infiltration of *N. benthamiana* leaves, the fluorescence emitted by pHGFP is recorded after an excitation wavelength of 405 nm and 488 nm, to establish a 405/488 fluorescence intensity ratio, indicative of pH differences. The discrimination between the apoplastic and cytosolic 405/488 ratio was made possible by the use of the following constructs. The apoplastic membrane pH sensor pHGFP-PM-Apo resulted from the fusion of pHGFP with the TMD of the PM-localized protein TM23 (Brandizzi et al., 2002), and the cytosolic membrane pH sensor pHGFP-PM-Cyto corresponded to the fusion of pHGFP with the C-terminal farnesylation sequence of Ras which is anchored to the PM (Martinière et al., 2018).

As expected, the 405/488 nm fluorescence ratio measured in *N. benthamiana* cells was higher for the pHGFP-PM-Cyto (median=2.2) when compared to that for pHGFP-PM-Apo (median=1.3), revealing the higher pH of the cytosolic compartment than that of apoplast (**Figure 1C**). The 405/488 nm fluorescence ratio was then measured in cells transformed with FW2.2 fused with the pHGFP either at its N-terminal or C-terminal end. It was shown to be very close to the fluorescence ratio measured with the pHGFP-PM-Apo (median=1.3), thus demonstrating unequivocally that the N- and C-terminal parts of FW2.2 are facing the apoplast (**Figure 1B**).

### FW2.2 is enriched at plasmodesmata

To go deeper into the study of the FW2.2 subcellular localization, we generated stable transgenic lines expressing FW2.2 fused to YFP at its C-terminal end under the control of the Cauliflower Mosaic Virus (CaMV) 35S promoter (referred to as *35S::FW2.2-YFP* plants), in the cultivated tomato variety Ailsa Craig (AC). In these plants, the emitted fluorescence associated to YFP was highly detectable in roots and leaves, and in reproductive organs, namely flowers and fruits (**Supplemental Figure 1A**). The localization of FW2.2-YFP at the PM was confirmed in all tissues investigated, namely in roots and fruit pericarp (**Figure 2A**), according to a pattern of punctate spots at the cell periphery, suggesting that FW2.2-YFP was enriched at nanodomains as observed previously for the soybean ortholog GmFWL1 (Qiao et al., 2017). The same tissue preparations were then stained with aniline blue (AB) to reveal callose deposition, as a marker of PD. The fluorescent dots revealing FW2.2-YFP co-localised with AB staining, at pit field junctions, as shown by the overlapping signal intensity plots (**Figure 2A**), thus indicating a localization at PD. It is noteworthy that the localization of FW2.2 at PD was independent from the position of YFP at the C-terminal or N-terminal end of the protein, since we obtained similar results using a *35S::YFP-FW2.2* construct (**Supplemental Figure 1B**). The enrichment of FW2.2 at PD was quantified by measuring the plasmodesmata enrichment ratio, named ‘PD index’, corresponding to the FW2.2-YFP fluorescence intensity at PD *vs* that at the cell periphery, as previously described (Brault et al., 2019; Grison et al., 2019). As a control, root and fruit pericarp tissues from WT plants were stained with FM4.64, a PM-specific dye (Bolte et al., 2004), together with AB to measure the PD index. While the PD index in controls was equal to 1 regardless of the tissue tested, a high PD-index ranging from 1.7 to 1.9 was measured in root and pericarp cells of *35S::FW2.2-YFP* plants, (**Figure 2B**), thus demonstrating that FW2.2 was enriched at PD.

**Figure 2.**
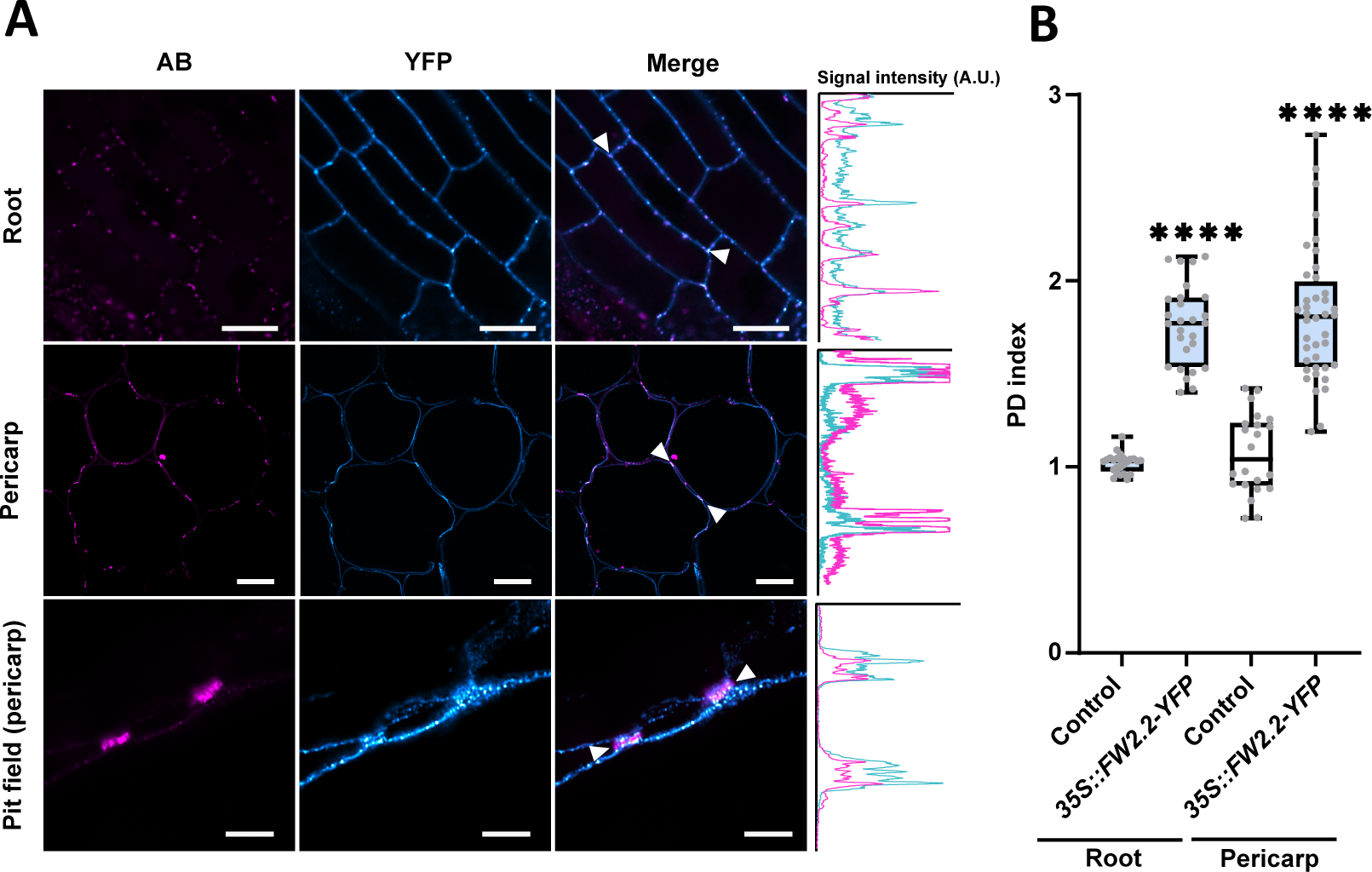
FW2.2 is enriched at PD. **(A)** Confocal microscope observations of FW2.2-YFP localization in roots, pericarp and pit field junctions in pericarp cells from *35S::FW2.2-YFP* plants. Scale bar for root and pericarp = 10 µm. Scale bar for pit field = 5µm. Intensity plots delineated by the two white arrowheads are shown for each co-localisation pattern. A.U. = Arbitrary unit. **(B)** PD index for FW2.2 in roots and pericarp tissue of *35S::FW2.2-YFP* plants compared to WT. n>20. Statistical analysis: Student’s t-test. \*\*\*\**P* < 0.0001.

### The overexpression of FW2.2 in leaves enhances cell-to-cell diffusion capacity

Since FW2.2 localizes at PD, we hypothesized that it could contribute to a function associated to cell-to-cell communication. To test this hypothesis, a new set of gain-of-function plants were generated in the tomato cultivar AC, as to overexpress *FW2.2* constitutively and ectopically, under the control the 35S promoter (referred to as *35S::FW2.2*). Three lines were selected with medium-(2-fold more) to very high levels (50-fold more) of *FW2.2* overexpression in 5 DPA fruits, a stage when the endogenous *FW2.2* expression is at its maximum (**Supplemental Figure 2**). In parallel, loss-of-function plants were generated using the CRISPR/Cas9 technology. To knock out *FW2.2*, two single-guide RNAs (sgRNAs) were designed as close as possible to the start codon of the coding sequence to create a frameshift or an early stop codon resulting in a dysfunctional FW2.2 protein in which the PLAC8 domain is missing (**Supplemental Figure 3**). Three different homozygous lines were selected, and referred to as *CR-fw2.2* hereafter.

In all three independent *35S::FW2.2* overexpressing lines, a significant reduction in mean leaf surface was observed, from 33% to 42% compared to that in WT (**Figure 3A**). This reduction in leaf surface was not due to any alteration of cell size, as the leaf epidermal cell density, used as a proxy for cell size, was unaffected (**Figure 3B**). No growth-related phenotype was observed in leaves of *CR-fw2.2* plants, which was expected as *FW2.2* is not naturally expressed in leaves (**Supplemental Figure 2B**).

**Figure 3.**
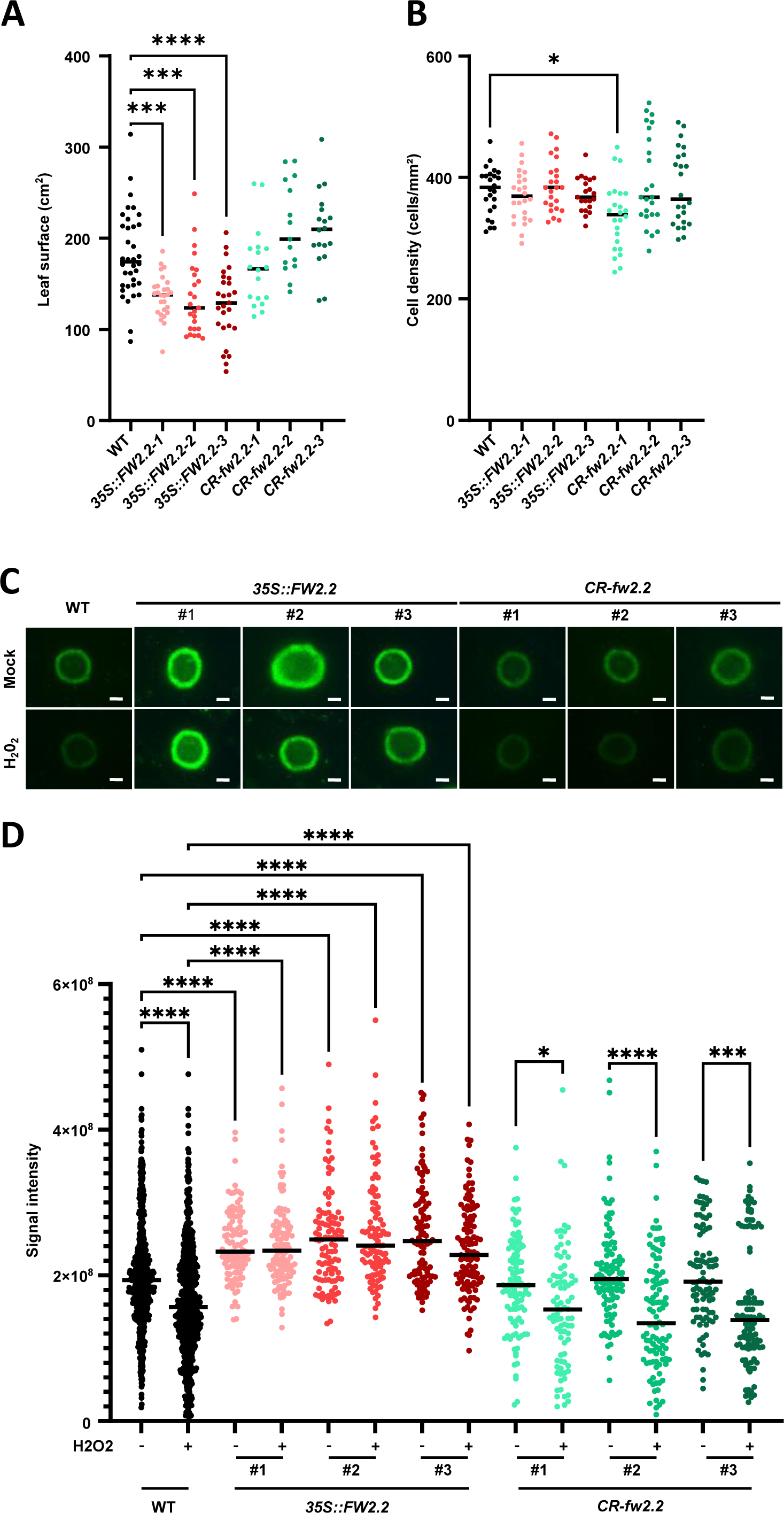
The overexpression of FW2.2 enhances cell-to-cell diffusion in leaves. **(A)** Determination of the mean mature leaf surface in WT, *35S::FW2.2* and *CR-fw2.2* lines. **(B)** Determination of the cell density in leaves from WT, *35S::FW2.2* and *CR-fw2.2* lines. **(C)** DANS assays using leaves from WT, *35S::FW2.2* and *CR-fw2.2* lines with or without H_2_0_2_ treatment. Scale bar = 500 µm. **(D)** Quantification of the CF signal intensity in WT, *35S::FW2.2* and *CR-fw2.2* lines with or without H_2_0_2_ treatment. Statistical analysis: Kruskal–Wallis test with post hoc Dunn multiple comparison test. \**P* <0.05; \*\*\**P* <0.001; \*\*\*\**P* <0.0001.

We next investigated whether the overexpression of *FW2.2* in leaves could affect the permeability of PD, and consequently the cell-to-cell communication. The PD permeability in WT, *35S::FW2.2* and *CR-fw2.2* lines was compared by performing “Drop-ANd-See” (DANS) quantitative assays (Cui et al., 2015), using the membrane-permeable, non-fluorescent dye Carboxy-Fluorescein DiAcetate (CFDA). DANS assays are based on the ability of cell to uptake CFDA rapidly; intracellular esterases then cleave CFDA into fluorescent but membrane-impermeable Carboxy-Fluorescein (CF), and CF diffuses symplastically into the neighbouring cells only via PD. To our knowledge, the use of this technique has never been reported in tomato. We first checked that DANS assays are functional in tomato using leaflets of 4 weeks-old plants (**Supplemental Figure 4A**).

In Arabidopsis, a pre-treatment with 10 mM H_2_0_2_ alters PD permeability through an increase in callose deposition (Cui and Lee, 2016). Such an effect was also observed in tomato WT leaves, as revealed by the reduction in CF signal intensity compared to mock-treated leaves, thus indicating a decrease in PD permeability affecting the cell-to-cell movement of CF in tomato leaves (**Figure 3C-D**). We then examined whether gain- or loss-of-function of *FW2.2* alters cell-to-cell communication. The CF signal intensity was increased (from 20 to 30%) in all overexpressing *35S::FW2.2* lines compared to that in WT, suggesting an increased PD permeability (**Figure 3C-D**). Interestingly, the H_2_0_2_ treatment which increases callose deposition in WT and thereby decreases PD permeability, had no effect on the *35S::FW2.2* lines, compared to the mock treatment. Hence, not only the overexpression of *FW2.2* in leaves increased PD permeability, but it also inhibited the negative effects of H_2_0_2_ on it. On the contrary, the CF signal intensity in *CR-fw2.2* lines was similar to that in WT (**Figure 3C-D**), showing no difference in CF diffusion, which suggests that the PD permeability was not affected. This absence of effects on PD permeability in *CR-fw2.2* lines can be explained by the absence of endogenous *FW2.2* expression in leaves, as mentioned above. It also correlates with the absence of any alteration in epidermal cell size in *35S::FW2.2* and *CR-fw2.2* lines (**Supplemental Figure 4B**). Therefore, the observed difference in CF diffusion was the result of the overexpression of *FW2.2* in tomato leaves, which induced a modification in the cell-to-cell communication status, as revealed by the altered PD permeability.

### FW2.2 affects the callose deposition at PD in leaves

A key mechanism for the regulation of PD aperture, and therefore for intercellular flux of signalling molecules, involves the accumulation of the cell wall polysaccharide callose at the neck regions of PD (Amsbury et al., 2018). To verify whether the increase in cell-to-cell diffusion mediated by the overexpression of *FW2.2* was due to a modified level of callose accumulation, the levels of callose at PD were measured in leaves from WT, *35S::FW2.2* and *CR-fw2.2* plants, following a pre-treatment with or without H_2_O_2_. The levels of callose were quantified by immunofluorescence labelling using a callose-specific antibody as illustrated for WT in **Figure 4A**, and the signal intensity was subsequently quantified as a proxy of callose deposition at PD (**Figure 4B**), as commonly used (Grison et al., 2019; Platre et al., 2022; Wang et al., 2023). Compared to control conditions (mock treatment), the signal intensity for callose in WT leaves treated with H_2_O_2_ was increased, correlating with DANS assays showing decreased cell-cell communication. The immunofluorescence intensity in the *35S::FW2.2* leaves was decreased when compared to that in WT, indicating that less callose was deposited, in the absence of any alteration in cell size and leaf thickness as verified before (**Figure 3B** and **Supplemental Figure 4**). In response to H_2_O_2_, the levels of callose deposition in *35S::FW2.2* leaves also increased, but to a much lower extent than in WT (**Figure 4B**). On the contrary, the levels of callose deposition in *CR-fw2.2* leaves with or without H_2_O_2_ were highly similar to that in WT, in accordance with the absence of phenotype when *FW2.2* is mutated (**Figure 3**).

**Figure 4.**
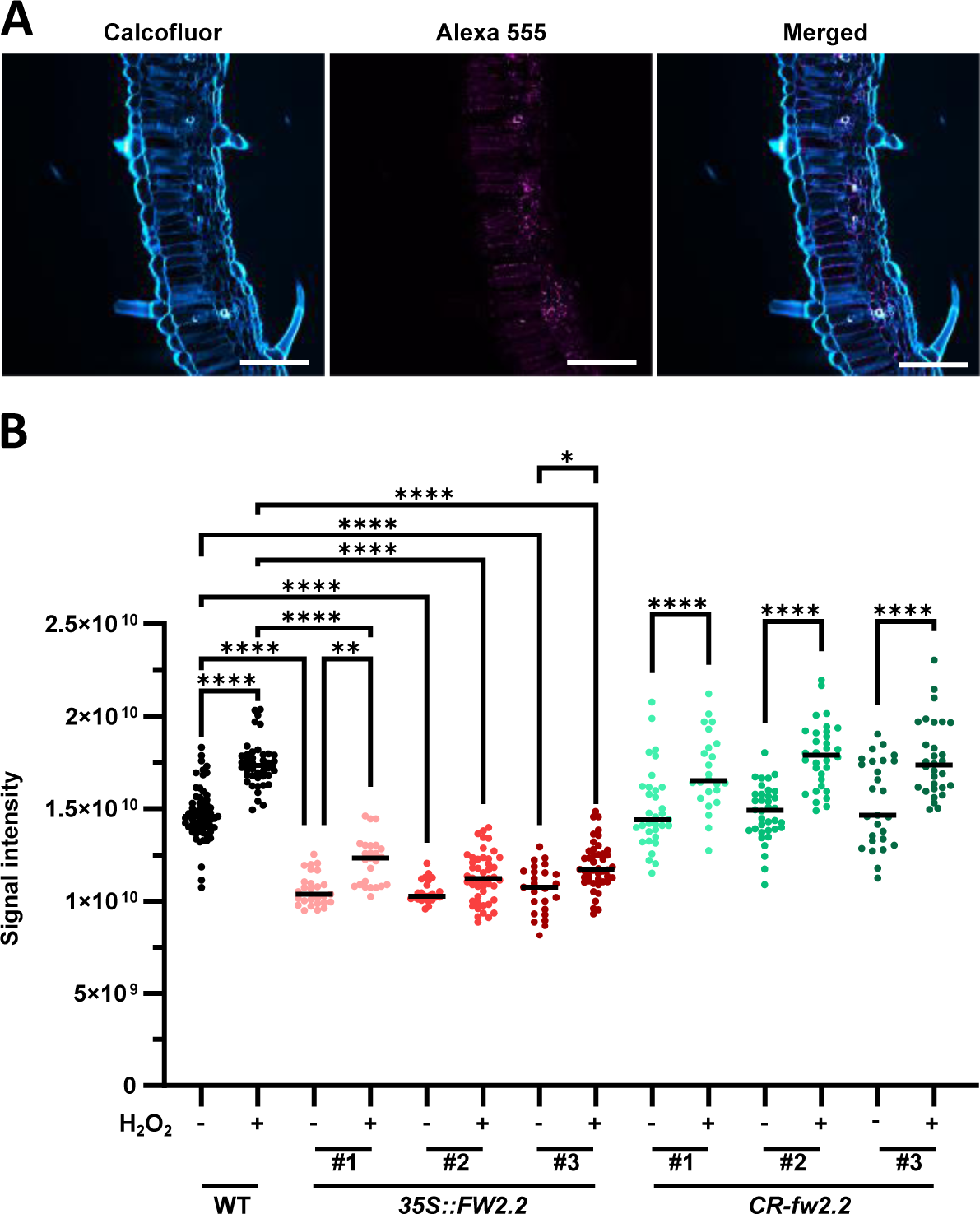
The overexpression of FW2.2 alters callose deposition in leaves. **(A)** Immuno-labeling of callose in leaves of WT plants. Scale bar = 100 µm. **(B)** Quantification of callose deposition in WT, *35S::FW2.2* and *CR-fw2.2* lines. The signal intensity for callose deposition is integrated to the pixel surface measured. Statistical analysis: Kruskal–Wallis test with post hoc Dunn multiple comparison test. \*\**P* <0.01; \*\*\**P* <0.001; \*\*\*\**P* <0.0001. n>20.

These results clearly indicated that FW2.2 alters the process of callose deposition at PD.

### FW2.2 regulates negatively callose deposition at PD in fruit pericarp

Since FW2.2 was found as a major regulator of fruit weight, we next examined whether the misexpression of *FW2.2* would affect the level of callose deposition at PD in fruit pericarp tissue.

At a macroscopic level, among the three selected overexpressing lines, a significant reduction in mean fruit weight was observed for the *35S::FW2.2-1* and *35S::FW2.2-3* lines (according to an average increase of 19.6% and 11.3% respectively) (**Figure 5A**). The mean fruit weight in the three *CR-fw2.2* loss-of function plants was higher than that of the WT (7,2%, 7,1% et 6,3% respectively). However, these differences were not statistically significant, because of a high variability in fruit weight values, In addition, there was no modification in pericarp thickness in mature fruits from the three *35S::FW2.2* lines compared to WT fruits, while pericarp from *CR-fw2.2* fruits appeared thinner (**Figure 5B**). Related to fruit structure, fruits from gain- and loss-of-function plants were all affected for the number of locules to various degrees (**Figure 5C**). More fruits with less than 3 locules were encountered in the overexpressing *35S::FW2.2* lines, while fruits with 4 and even more locules were observed in *CR-fw2.2* lines, compared to WT fruits from the AC cultivar which usually contain 3 locules. This converse impact on the number of fruit locules in the gain- and loss-of-function plants suggests that cell divisions have been impacted in the floral meristem (FM) termination process, through the increased or repressed negative regulatory effect in *35S::FW2.2* or *CR-fw2.2* lines respectively.

**Figure 5.**
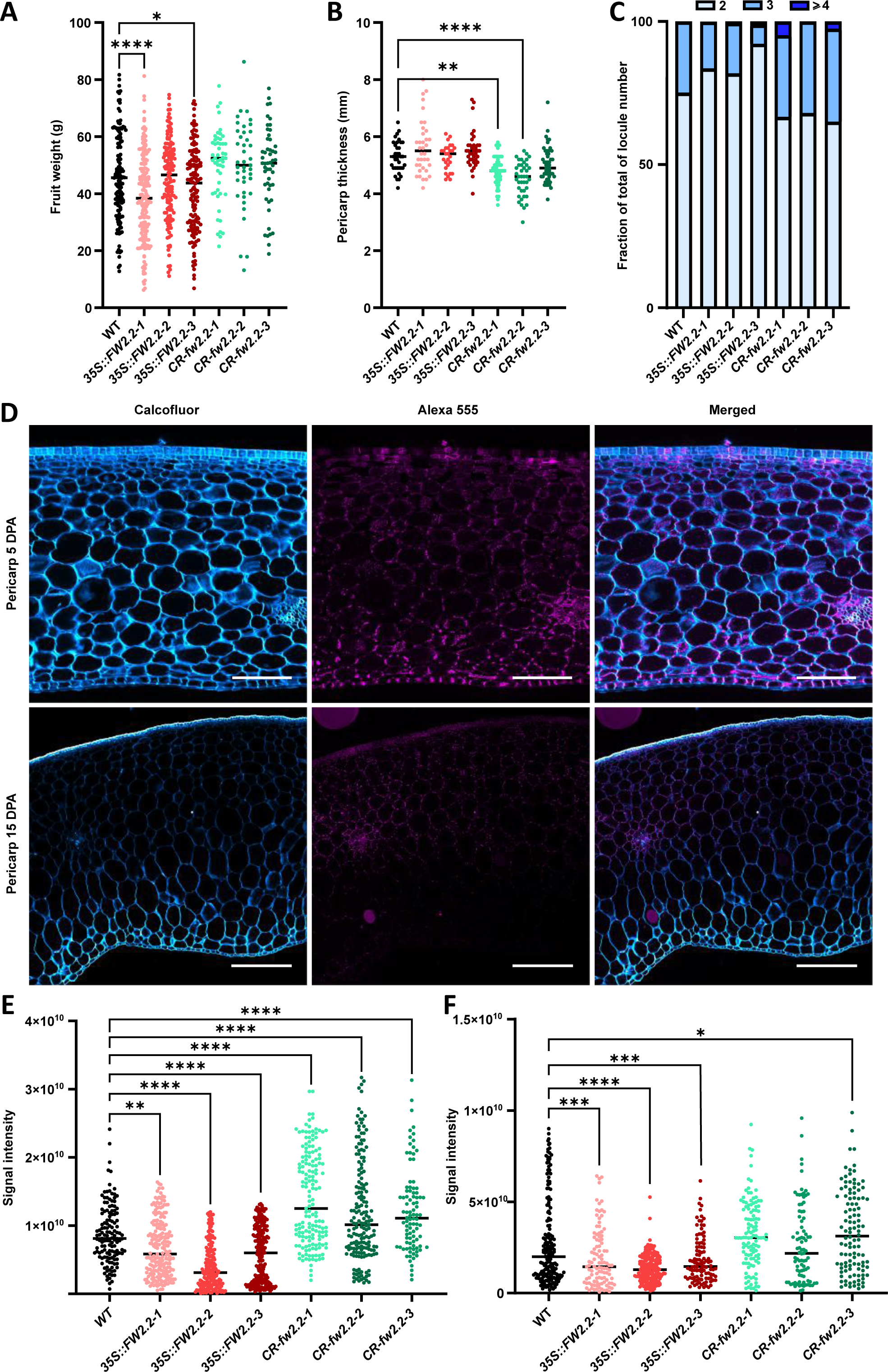
Callose deposition is altered at 5 and 15 DPA in fruit pericarp of *35S::FW2.2* and *CR-fw2.2* plants. **(A-C)** Phenotypic analysis of fruits (at breaker stage) from *35S::FW2.2* and *CR-fw2.2* plants compared to that of WT: Determination of the mean fruit weight (**A**); Determination of the pericarp thickness **(B)**; Determination of the number of fruit locules (**C**). **(D)** Immunolabeling of callose in 5 DPA (top) and 15 DPA (bottom) pericarp from WT fruits. Scale bar = 100 µm (top); 500 µm (bottom). **(E-F)** Level of callose deposition in WT, *35S::FW2.2* and *CR-fw2.2* lines at 5 (**E**) and 15 DPA (**F**). The signal intensity for callose deposition is integrated to the pixel surface measured. Statistical analysis: Kruskal–Wallis test with post hoc Dunn multiple comparison test. \**P* <0.05; \*\**P* <0.01; \*\*\**P* <0.001; \*\*\*\**P* <0.0001. n>80.

The level of callose deposition was then investigated on pericarp sections of fruits from the *35S::FW2.2* and *CR-fw2.2* plants harvested at 5 and 15 DPA (**Figure 5D**). These two different developmental stages were chosen because *FW2.2* is highly expressed in the pericarp of 5 DPA fruit and much less at 15 DPA (**Supplemental Figure 2B**). At both 5 and 15 DPA, the immunofluorescence signal intensity in the pericarp of *35S::FW2.2* fruits was decreased when compared to that in WT, indicating that the level of callose deposition was reduced (**Figure 5D-E**). On the contrary, the immunofluorescence signal intensity in the pericarp of *CR-fw2.2* fruits at both 5 and 15 DPA was increased significantly when compared to that in WT, thus revealing a higher level of callose deposition. Interestingly, the increase in callose deposition observed at 15 DPA in pericarp sections from *CR-fw2.2* fruits was less pronounced than at 5 DPA, and almost identical to that in WT. This can be explained by the very low expression of *FW2.2* in 15 DPA fruits (**Supplemental Figure 2B**), and thus the absence of any loss-of-function effect from the CRISPR-Cas9 construct on *FW2.2* at this developmental stage.

Cell perimeters were measured for all genotypes in all the different cell layers composing the fruit pericarp at 5 DPA, and in the mesocarp at 15 DPA, to ascertain that these differences in callose deposition was not due to any heterogeneity in cell size, and thus in the density of cell walls. The cell perimeter was comparable in all WT, *35S::FW2.2* and *CR-fw2.2* lines, with only slightly smaller values in some cases, especially in the internal part of the mesocarp (**Supplemental Figure 5**). Hence, the observed differences in callose deposition did originate from the effects of *FW2.2* gain- and loss-of-function, demonstrating that FW2.2 regulates negatively the process of callose deposition at PD within fruit pericarp.

### FW2.2 interacts physically with Callose Synthases

To go deeper into the functional and biochemical characterization of FW2.2, an *in vivo* approach using immunoprecipitation followed by tandem-mass spectrometry (IP-MS/MS) was performed to identify interacting protein partners of FW2.2 inside the pericarp from *35S::FW2.2-YFP* fruits harvested at 10 DPA. Since *FW2.2* is still expressed endogenously at this developmental stage, it was therefore expected that its natural interacting proteins would be present in the protein extracts. The IP-MS/MS experiment resulted in the identification of 662 proteins interacting with FW2.2, which were enriched in the *35S::FW2.2-YFP* sample when compared to WT (**Figure 6A**, **Supplemental Data Set 1**). To identify potential PD-localized candidates in relation with FW2.2 function, we compared this list with a tentative PD proteome from tomato made of a total of 400 proteins corresponding to the deduced orthologs of the 115 proteins constituting the refined PD proteome from Arabidopsis published by Brault *et al*. (2019). Seventeen proteins were found overlapping between the two proteomes (**Figure 6B**). Three distinct classes of proteins, all key regulators of cell-to-cell signalling in plants, represented almost two thirds of the identified proteins (**Figure 6C**): i) two proteins of the C2 calcium/lipid-binding phosphoribosyl transferase family (Solyc01g080430 and Solyc01g094410), belonging to the large family of multiple C2 domains and transmembrane region proteins (MCTP) (Brault et al., 2019); ii) three proteins of Leucine-Rich Repeat Receptor-Like kinases (LRR-RLKs) family (Solyc03g111670, Solyc06g082610 and Solyc05g052350) (Wei et al., 2015); iii) six different Callose Synthases (CalS), which were identified based on their phylogenetic proximity to Arabidopsis counterparts, namely SlCalS1 (Solyc01g006350), SlCalS3a (Solyc01g006370), SlCalS3b (Solyc01g073750), SlCalS9 (Solyc01g006360), SlCalS10a (Solyc03g111570) and SlCalS12 (Solyc07g053980) (**Supplemental Figure 6A**). The preferential interaction of FW2.2 with Callose synthases in 10 DPA fruits was thus fully relevant with its aforementioned role in regulating callose deposition at PD in the pericarp. RT-qPCR analyses confirmed that these 6 *CalS* genes were expressed in WT fruit pericarp at 10 DPA (**Supplemental Figure 6B)**. In addition, there was no significant change in the expression level of the 6 *CalS* genes in tomato leaves and fruits at 5 and 15 DPA from the *FW2.2* loss- and gain-of-function plants except for SlCalS12 (Solyc07g053980) whose expression was lower in leaves and 5 DPA fruits of *35S::FW2.2* and higher in 5 DPA fruits of *CR-fw2.2* (**Supplemental Figure 7**). Therefore, the preferential interaction between FW2.2 and the six CalS proteins is not related to an increase in *CalS* gene expression in the *35S::FW2.2-YFP* plants, and ultimately to an increased translation, but to the endogenous level of CalS in the protein extracts of 10 DPA fruits used for the IP experiment.

**Figure 6.**
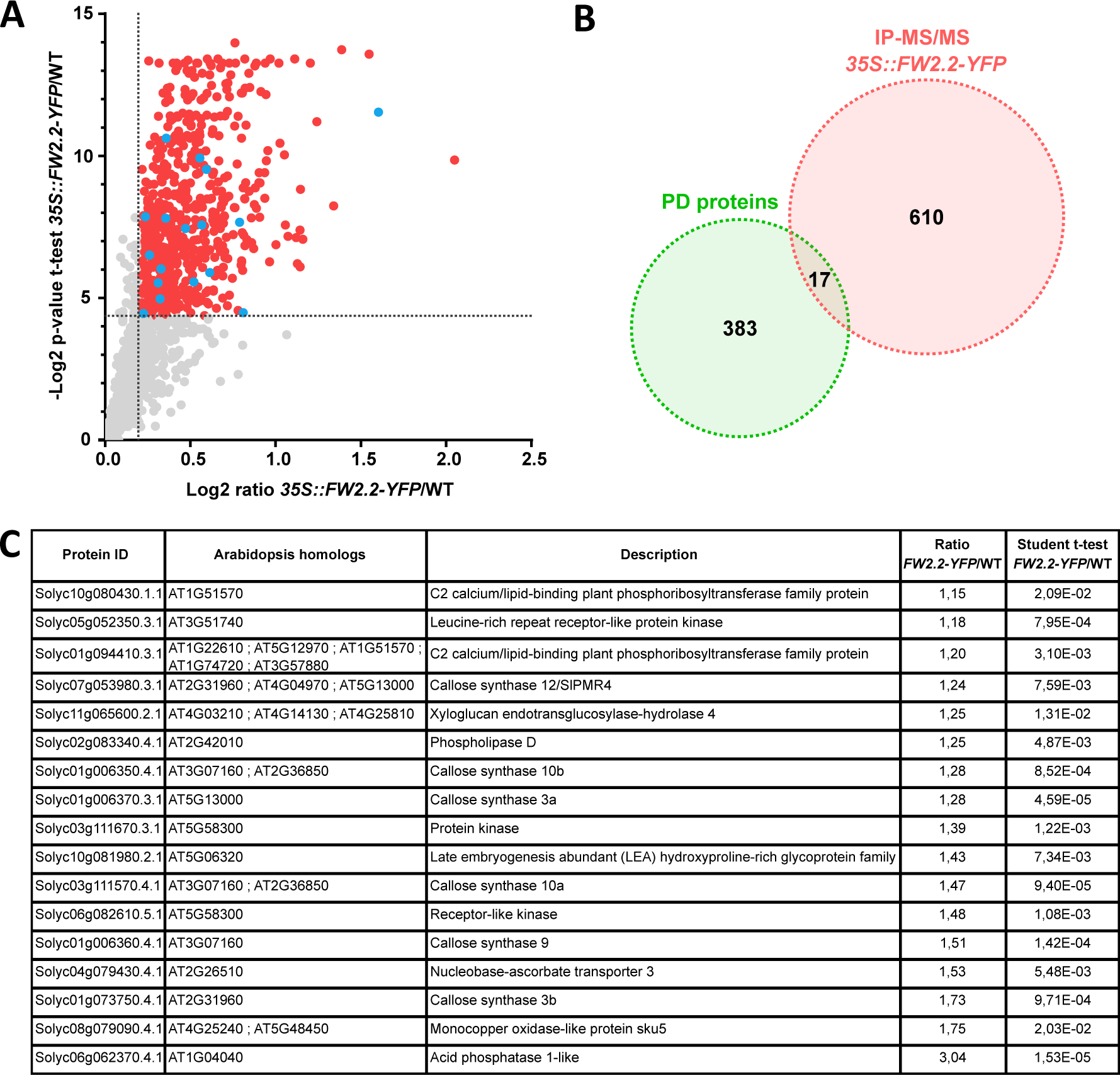
FW2.2 physically interacts with several PD localized protein including callose synthases. **(A)** Dot plots showing enriched proteins in *35S::FW2.2-YFP* IP-MS/MS experiments in 10 DPA pericarp. Red dot indicates significantly enriched protein (based on a Student’s t-test with Benjamini-Hochberg correction *P* < 0.05 and an enrichment ratio > 1.15). Blue dots indicate proteins found in the PD proteome. **(B)** Venn diagram showing the overlap between the IP-MS/MS proteome and the PD proteome. Statistical analysis: Hypergeometric test *P*=0.0021. **(C)** List of plasmodesmata proteins detected in the IP-MS/ proteome.

These results thus support the functional role of FW2.2 on PD permeability and cell-to-cell communication, via an interaction with Callose synthases, which may potentially modulate their catalytic activity.

## DISCUSSION

FW2.2 was the first gene underlying a QTL related to fruit size to be cloned in tomato (Frary et al., 2000). It is by far the major QTL of such type, as it accounts for as much as a 30% difference in fruit fresh weight between domesticated (large-fruited) tomatoes and their wild (small-fruited) relatives (Frary et al., 2000; Grandillo et al., 1999). Most wild -small fruited-tomatoes (if not all) possess ‘small-fruit’ alleles; conversely all domesticated/cultivated -large fruited-tomatoes possess ‘large-fruit’ alleles (Bianca et al., 2015). Comparative sequence analysis of FW2.2 from the large- and small-fruited alleles indicated that the *FW2.2* effects on fruit size do not originate from differences in the sequence and structure of the protein, but rather from the timing of its transcription (heterochronic changes) and the overall quantity of transcripts in the fruit (Cong et al., 2002). The ‘large-fruit’ allele is rapidly transcribed to reach a peak of expression around 5 DPA, whereas the ‘small-fruit’ allele is transcribed more slowly and displays its maximum of expression nearly a week later (12 to 15 DPA), reaching almost twice the mRNA level observed in large-fruit allele (Cong et al., 2002). Since this difference in timing of expression was found inversely correlated to the mitotic activity, FW2.2 was defined as a negative regulator of cell divisions in pre-anthesis ovary and developing fruit, thus modulating final fruit size (Frary et al., 2000; Cong et al., 2002). Such a function in regulating organ size by modulating cell number was found conserved for many other plant orthologues of FW2.2 (Beauchet et al., 2021), which led to the attribution of the CELL NUMBER REGULATOR (CNR) protein family name (Guo et al., 2010). Members of the CNR protein family are targeted to the PM, due to the presence of the PLAC8 domain (Beauchet et al., 2021). However, the precise biological function and mechanism of action of membrane-embedded FW2.2 and CNRs in controlling organ size via the regulation of cell divisions remained totally elusive so far.

### FW2.2 regulates cell-to-cell diffusion by modulating callose deposition at plasmodesmata

It was long known that FW2.2 is a plasma membrane-located protein (Cong and Tanksley, 2006). Using transient expression in tobacco leaves and stable transformants in the tomato AC cultivar, we confirmed this PM localization for FW2.2 (**Figures 1-2**). The topology of FW2.2 within the PM was established and revealed that the N- and C-terminal regions are extracellular, thus facing the apoplast, while the protein loop in-between the two TMDs is cytoplasmic (**Figure 1**). These results were in full agreement with a topological model predicted for PfCNR1, the FW2.2 putative orthologue from *Physalis floridana*, which displays a high degree of homology with FW2.2 (Li and He, 2015). More importantly, we demonstrated unequivocally that FW2.2 is enriched at PD (**Figure 2**) and participates in cell-to-cell communication mechanisms via the regulation of PD permeability (**Figures 3**).

This localization at PD is most probably functionally conserved with other members of the CNR family. Indeed, the localization of the soybean GmFWL1 protein was described as associated to membrane microdomains (Qiao et al., 2017), according to a punctate pattern very similar to what we observed for FW2.2 in tomato (**Figure 2**). It is thus highly probable that GmFWL1 also localizes at PD. The closest homolog of FW2.2 in Arabidopsis, namely AtPRC2, belongs to the PD proteome established by Brault et al. (2019), together with well-established PD proteins, and presents a ∼50- to 100-fold enrichment at PD compared to the PM, total protein, microsomal or cell wall fraction.

PD make the connection between adjacent cells to enable the diffusion of mobile signalling molecules (Wu and Gallagher, 2011). Using DANS assays, we demonstrated that FW2.2 is involved in cell-to-cell diffusion mechanisms and contributes to increase PD permeability (**Figure 3**). The permeability and thus the aperture of PD are mechanically regulated by the extent of deposited callose at the neck of PD (Amsbury et al., 2018). The increase in PD permeability mediated by FW2.2 occurs via a modification in the level of callose deposition, as FW2.2 regulates negatively its accumulation (**Figures 4-5**). The level of callose deposition is a highly regulated process involving two antagonistic enzymes, Callose Synthases and β-1,3-glucanases (Chen and Kim, 2009). Callose deposition is enhanced according to two main signalling pathways, one Reactive Oxygen Species (ROS)-dependent and the other one salicylic acid (SA)-dependent, which both induce the expression of receptor proteins such as PDLP5 that participate with Callose Synthase proteins in the regulation of PD permeability (Cui and Lee, 2016; Amsbury et al., 2018; Tee et al., 2022). The expected decrease in PD permeability under H_2_0_2_ stress was not observed when FW2.2 is overexpressed, suggesting that FW2.2 play a role in the ROS-dependent pathway. Whether FW2.2 plays also a role in the SA-dependent pathway to regulate PD permeability remains to be determined.

### FW2.2 interacts physically with Callose Synthases to modify their activity

A proteomics approach using IP-MS/MS revealed that FW2.2 interacts with different Callose Synthases: SlCalS1, SlCalS3a, SlCalS3b, SlCalS9, SlCalS10 and SlCalS12 (**Figure 6**). Interestingly, all these tomato proteins are the orthologs of Arabidopsis CalS known to contribute to callose homeostasis at PD, thereby regulating the permeability of PD and consequently the symplastic molecular exchanges between neighboring cells (Saatian et al., 2023; Usak et al., 2023). It is noteworthy that among the 178 proteins found to interact with GmFWL1, three distinct callose synthases, namely CalS5 (Glyma13g31310), CalS8 (Glyma04g36710) and CalS10 (Glyma10g44150) were also identified following the co-immunoprecipitation assays (Qiao et al., 2017). This observation not only suggests that GmFWL1 is probably located at PD as well, but also that the interaction between FW2.2 and CNRs with proteins involved in the biosynthesis of callose and the metabolic process of callose deposition at PD seems to be a conserved feature for the balance between synthesis and degradation of callose at PD, and suggests that CNRs regulate negatively the activity of Callose Synthases.

CalS are very large proteins (more than 1900 aa) which possess multiple transmembrane spanning domains arranged in two regions delineating a cytoplasmic hydrophobic loop. This hydrophilic loop harbors the catalytic domain of the active CalS complex where UDP-Glucose transferase (UGT1) and Sucrose Synthase (Susy) may interact to provide substrates for callose synthesis (Verma and Hong, 2001). As revealed by the topological analysis of FW2.2, the protein sequence between the two TMDs corresponds to a cytoplasmic/intracellular region (**Figure 1**). It is likely that this cytoplasmic region is involved in the interaction with CalS proteins and other putative interactors, as also shown for PfCNR1 (Li and He, 2017).

The activity of PD-associated Callose Synthases is of prime importance in numerous developmental processes, such as in response to biotic and abiotic stress, organ and tissue patterning, cell differentiation, phloem transport, and cell division via the formation of the cell plate at cytokinesis (Amsbury et al. 2018; Wu et al., 2018; Usak et al., 2023). In Arabidopsis, AtCalS1 and AtCalS10 localize at the nascent cell plate where they synthesize callose as the first and fundamental polysaccharide component of the nascent cell plate, and AtCalS9 is essential for the proper commitment to mitosis during male gametogenesis (Usak et al., 2023). Again, orthologs for these three CalS were found to interact with FW2.2 in tomato. Interestingly, the CRR1 protein from rice encodes a CalS which is essential for ovary growth following fertilization (Song et al., 2016). The loss-of-function of *CRR1* induces a disordered patterning of vascular cells in the ovaries of the mutant, with aberrant cell wall formation and reduced callose deposition at PD. Furthermore, the cell number inside the *crr1* ovaries is reduced when compared to the WT, establishing a link with callose synthesis and deposition, symplastic pathway via PD and control of cell division during ovary development.

### How to reconcile a function of FW2.2 in cell-to-cell communication, cell cycle- and fruit growth regulation?

As FW2.2 was described as a negative regulator of cell division during early fruit development, which ultimately impacts fruit growth (Cong et al., 2002), it would have been expected that a loss-function of FW2.2 results in increased cell divisions and possibly larger organs (including fruits), and conversely that the ectopic overexpression of FW2.2 reduces mitotic activities and results in smaller organs. This latter effect could be observed at least in leaves from *35S::FW2.2* overexpressing lines (**Figure 3**), *i.e.* in organs where FW2.2 is not naturally expressed (**Supplemental Figure 2B**). Since the reduction in leaf growth was unrelated to any modification in cell size, this suggests that cell divisions were reduced under the effects of FW2.2 overexpression. In two out of three gain-of-function lines, we could also observe such a phenotype of reduced size for fruits although limited in extent (**Figure 5**).

These results are puzzling since genetics studies showed that the *fw2.2* QTL accounts for 22% to 47% of fruit mass variation when cultivated tomato cultivars are crossed with the wild species *Solanum pimpinellifolium* or *Solanum pennellii* (Alpert et al., 1995; Lippman and Tanksley, 2001; van der Knaap and Tanksley, 2003). Nevertheless, the literature is still devoid of any functional characterization of *FW2.2* in cultivated tomato plants, albeit the gene was discovered and cloned more than 20 years ago. This is most probably the result of a lack of phenotypes when *FW2.2* is artificially deregulated in transgenic fruits. For instance, Zsögön et al. (2018) aimed at introducing by CRISPR-Cas9 engineering, yield and productivity traits from modern (‘large-fruited’) tomato cultivars into the wild (‘small-fruited’) tomato *Solanum pimpinellifolium*. Among the six traits studied, these authors selected the *FW2.2* locus for fruit weight, and produced several mutants with deletions disrupting FW2.2. However, none of them induced any change in fruit size in T2 lines compared to *S. pimpinellifolium* WT, despite the mutations (Zsögön et al., 2018). These results corroborate the functional analysis reported herein in *S. lycopersicum* cv AC, when *FW2.2* was mutated in the *CR-fw2.2* loss-of-function plants (**Figure 5**). Hence, the ectopic and constitutive expression of *FW2.2* driven by the 35S promoter, definitely outside its natural timeframe and territorial regulation, and its loss of function did not impact fruit development, which probably obeys to precise changes in *FW2.2* spatio-temporal expression, according to the heterochronic regulation of expression described for the original *fw2.2* mutation (Cong et al., 2002). To cope with this difficulty, we developed an ‘allele swapping’ complementation strategy (**Supplemental Figure 8**). This strategy aimed at generating transgenic plants in which the ‘large-fruit‘-allele promoter from *S. lycopersicum* cv. AC is used to govern the expression of *FW2.2* in a ‘small-fruit’ background, namely the wild tomato *S. pimpinellifolium* (Pi). Conversely, we used the ‘small-fruit‘-allele promoter from *S. pimpinellifolium* to govern the expression of *FW2.2* in the ‘large-fruit’ AC background. Although we succeeded in the expected allele expression swapping according to the right spatio-temporal expression governed by each of the promoters, we failed to produce any fruit weight phenotypes in the complemented *S. pimpinellifolium* and *S. lycopersicum* cv. AC transgenic lines compared to WT plants. Therefore, the effects of *FW2.2* on fruit size obeys probably to a subtler regulation than the sole quantity of transcripts and availability of the protein. In addition, we cannot exclude that this lack of tangible phenotype may be related to gene redundancy within the *CNR/FWL* family, as 11 genes paralogous to *FW2.2* have been reported (Beauchet et al., 2021).

Despite the lack of consistent phenotypes when *FW2.2* is misexpressed, the functionality of the protein itself within its cellular and protein environment may be of prime importance. The discovery of the FW2.2 function in cell-to-cell communication via PD thus raises the question of its link with the regulation of cell division, and subsequent fruit size control. By impairing callose deposition and thus maintaining PD aperture, FW2.2 may contribute to facilitate the diffusion of signalling molecules whose nature is still unknown. As reviewed by Han et al. (2014b), TFs are well characterized examples of such signalling molecules that could play an important part in the determination of fruit size. Recently, it was shown that a cold stress increases callose accumulation in the FM of tomato plants, resulting in impaired feedback loops which regulate the activity of WUSCHEL (WUS) (Wu et al., 2023). The TF WUS specifies the maintenance of stem cell activity in FM, and therefore a deregulation of WUS activity impacts the number of carpel primordia, and ultimately the number of locules inside the fruit. As a consequence of cold stress, *CalS* genes are induced and promote the callose deposition in the FM, which blocks the PD-mediated symplastic connection and alters the cell-to-cell movement of WUS which no longer can exert its negative regulatory action on *CLAVATA3* and *AGAMOUS*. As a result, the activity of *WUS* is not terminated in due time, which leads to increased cell divisions in the FM producing extra carpels and locules during fruit organogenesis (Wu et al., 2023). Interestingly, we observed similar trends in our transgenics plants: a higher number of locules resulting from increased cell divisions in FM in loss-of-function CR-*fw2.2* lines, as callose deposition was increased, and the opposite effects in gain-of-function *35::FW2.2* lines (**Figure 5C**). FM termination requires the repression of WUS via a transcriptional repressor complex, involving the INHIBITOR OF MERISTEM ACTIVITY (IMA) protein (Bollier et al., 2018), which was described as a negative regulator of cell divisions. In particular, the overexpression of *IMA* leads to smaller fruits, while its repression enlarges the FM and leads to an increase in the locule number (Sicard et al., 2008). IMA and its transcriptional regulatory machinery may thus represent such signalling molecules whose diffusion across PD may be influenced by FW2.2 to determine fruit size, as *FW2.2* is expressed as early as in carpels of pre-anthesis floral buds (Frary et al., 2000), and preferentially expressed in the FM than in vegetative meristems (Park et al., 2012).

So far, direct evidences for the symplastic movements via PD of cell cycle regulators have not been reported. However, Weinl et al., (2005) showed that Cyclin-Dependent Kinase (CDK)-specific inhibitors called Kip-Related Proteins (KRPs) can act non-cell-autonomously, as to regulate cell division and growth pattern in leaf epidermis. During tomato fruit development, KRPs are key players in the regulation of cell cycle, and the commitment to endoreduplication which drives ploidy-dependent fruit growth (Bisbis et al., 2006; Nafati et al., 2011; Tourdot et al., 2023). Whether the negative regulation on cell division exerted by FW2.2 in fruit growth goes through the inactivation of CDK/Cyclin activities via the traffic of KRPs from cell to cell across the pericarp remains an exciting matter of investigation. Recently, Ruan et al. (2020) reported that OsCNR1, encoded by the underlying gene of a major QTL for grain width and weight in rice, is able to interact with OsKRP1 in the cell membrane. Therefore, this remarkable finding provided the first evidence of a direct link between a CNR protein controlling organ size and a well-established cell cycle regulator inhibiting cell division. Whether this applies to FW2.2 for the regulation of cell cycle during early fruit development is a challenge for future research as to unravel definitely the function of FW2.2 in the control of fruit size/weight in tomato. Then, the lack of phenotypes observed in our in planta functional analysis may not be only related to the proper spatio-temporal expression of *FW2.2*, but also to the protein environment itself and the spatio-temporal availability of these putative signaling molecules.

How PD-mediated symplastic signalling affects fruit growth is still poorly understood. By demonstrating that FW2.2 contributes to the spatio-temporal regulation of callose deposition dynamics via regulating the CalS activity, we here provide an important breakthrough for the identification of the molecular and cellular mode of action of FW2.2. Based on our data, we propose a model integrating FW2.2 in the regulation of PD aperture via the dynamics of callose deposition (**Figure 7**). We propose that FW2.2 interacts with CalS to regulate negatively its activity, thus impacting PD permeability and facilitating the cell-to-cell movement of mobile signalling molecules. A future challenge will be to identify the nature of such signalling molecules, which will provide a valuable insight into the molecular mechanisms underlying the complex regulation of organ size, especially fruits.

**Figure 7.**
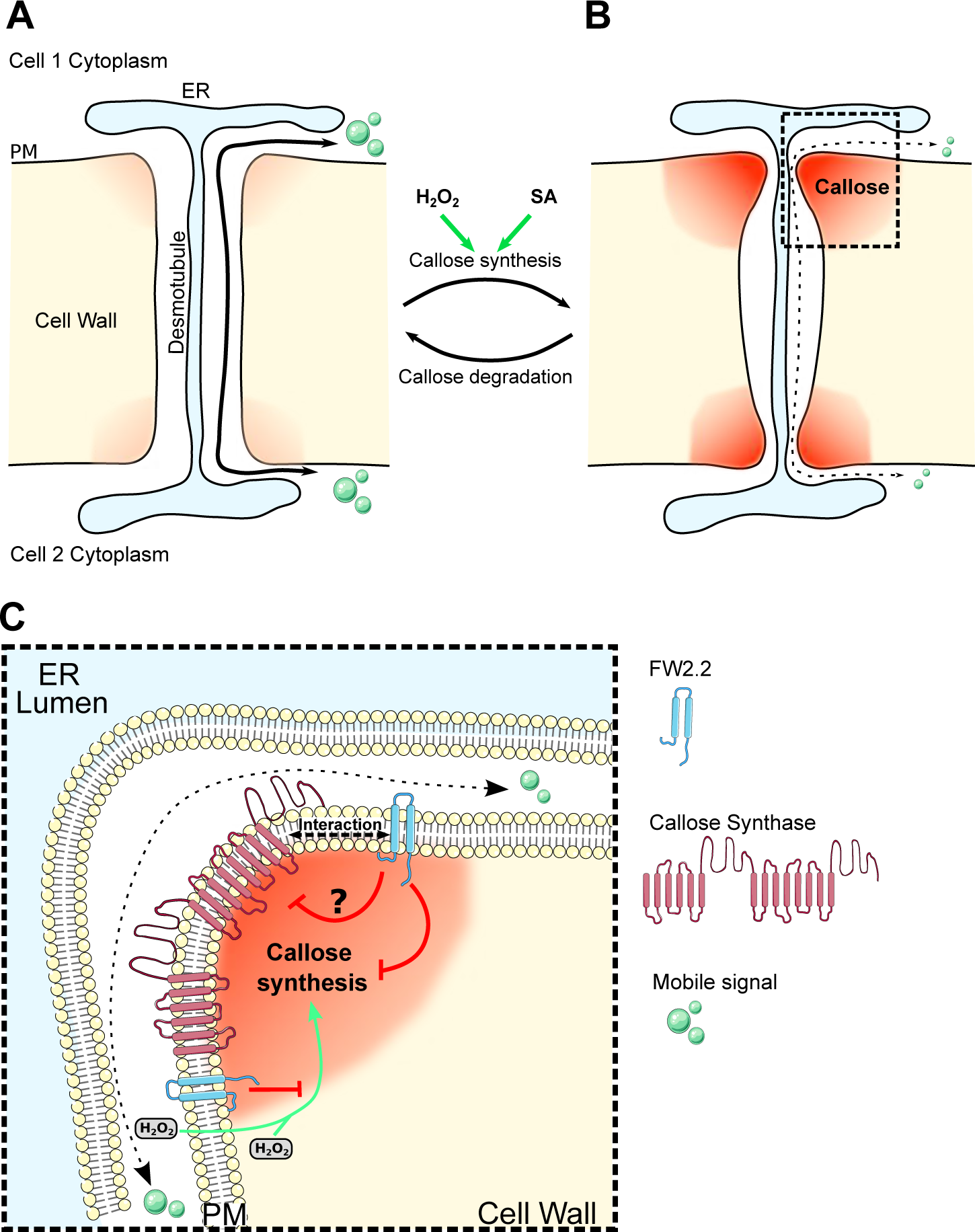
Model illustrating the function of FW2.2 in regulating callose synthesis at PD. **(A)** Regulation of PD aperture by callose deposition at the neck region of PD. PD aperture is regulated by the turn-over of callose: (**A**) a low callose deposition enables PD opening and facilitates the traffic of signalling molecules; (**B**) callose deposition, enhanced by ROS (H_2_O_2_) and SA, restricts the aperture of PD and the size of signalling molecules passing through. (**C**) Molecular and cellular model for the regulation of Callose synthase activity by FW2.2 at PD.

## METHODS

### Plant materials and growth conditions

Tomato (*Solanum lycopersicum* cv. AC) and tobacco (*Nicotiana benthamiana*) plants were grown in soil in a greenhouse under the following conditions: 16 h day/8 h night cycle, using a set of 100 W warm white LED projectors providing an irradiance of 100 μmol m^-2^ s^-1^ at the level of canopy. The light spectrum was constituted by equivalent levels of blue irradiation (range 430–450 nm) and red irradiation (640–660nm). For in *vitro* culture, tomato seeds were sterilized for 10 min under agitation in a solution of 3.2% bleach. Seeds were then washed three times with sterile water and dried under a laminar flow hood. Seeds were sowed in Murashige and Skoog medium (1/4 MS) and transferred in a growth chamber under the following conditions: 16 h day/8 h night cycle, 22°C/20°C day/night, using white light (Osram L36 W/77 Fluora 1400 Im) providing 80 to 100 μE m^-2^ s^-1^ intensity light at the stirring plate.

### Vector constructs and plant transformation

Vectors for the overexpression of *FW2.2* in plants were generated using the Gateway® cloning system (Invitrogen, Carlsbad, CA, USA), following manufacturer’s instruction. The *FW2.2* full-length coding sequence was amplified from cDNAs prepared from tomato (cv. AC) fruits at 5 DPA using PrimeSTAR MAX DNA polymerase (TAKARA BIO Inc., Kusatsu, Japan) and primers including the attB sites (**Supplemental Table 1**). The resulting PCR products were cloned into the corresponding Gateway vectors described in **Supplemental Table 2**. For CRISPR/Cas9 mutagenesis, constructs were assembled using the Golden Gate cloning method (Weber et al., 2011). Two sgRNAs were designed at the 5’ end of the coding sequence of *FW2.2* using CRISPOR (Concordet and Haeussler, 2018) to generate a premature stop codon (**Supplemental Table 1**). Primers for creating the sgRNA were designed as follows: tgtggtctcaATTG-NNNNNNNN-gttttagagctagaaatagcaag as a forward primer containing the sgRNA, and tgtggtctCAAGCGTAATGCCAACTTTGTAC as a reverse primer. The sequences corresponding to the sgRNA were then PCR amplified using the two aforementioned primers, and cloned into the pSLQ1651-sgTelomere plasmid (Addgene #51024). *fw2.2*-sgRNA-1 and *fw2.2*-sgRNA-2 were fused to the Arabidopsis AtU6-26 promoter (Addgene #46968) by digestion-ligation reaction in plCH47751 (Addgene #48002) and plCH47761 (Addgene #48003) respectively. These two level 1 vectors were assembled with the Kanamycin resistance gene (pNOS::NPTII-OCST; Addgene #51144), the AtCas9 (2×35S::AtCAS9-OCST; Addgene #112079) and the linker pICH41780 (Addgene #48019) into the level 2 vector plCSL4723 (Kind gift from Dr Mark Youles, The Sainsbury Laboratory, Norwich, UK). Transgenic plants were generated by *Agrobacterium tumefaciens* (strain C58C1) mediated transformation using explants of tomato cotyledons as described (Swinnen et al., 2022).

### RNA extraction and RT-qPCR analysis

Total RNA was isolated from cotyledons, hypocotyls, shoot apical meristems, leaves, roots, flowers and pericarp tissues from fruits harvested at different developmental stages (5, 10, and 15 DPA), using TRIzol reagent (Invitrogen) in combination with RNeasy Plant Mini Kit (Qiagen) following the manufacturers’ instructions. RNase-free DNase (Qiagen) treatment was performed on each sample. Reverse transcription was performed using the iScript^TM^ cDNA Synthesis Kit (Bio-Rad, Hercules, CA). Real-time PCR was performed using Gotaq® qPCR mastermix (Promega, Madison, WI) and a CFX 96 real-time system (Bio-Rad). qPCR primers were designed with PerlPrimer software (Marshall, 2004) to overlap 2 exons in order to limit genomic DNA amplification (**Supplemental Table 1**) and amplify a 80 to 200 bp-long amplicon, with a Tm of 60°C. The transcript levels of the expressed genes were normalized to that of the housekeeping genes: *SlTUBULIN* (Solyc04g081490) in combination with *SlNUDK* (Solyc01g089970) for fruit samples, or with *SlEIF4α* (Solyc12g095990) for other tissue samples.

### Phenotypic characterization

Plants were cultivated randomly side-by-side with WT plants. Flowers were vibrated every day to ensure optimal self-pollination. Seven flowers per inflorescence were maintained to ensure proper development of fruit per inflorescence. Fruits from four to six plants of each genotype of two biological replicates were used to determine fruit weight, fruit size, locule number and pericarp thickness at the breaker stage of fruit development. Fruits were weighted and measured using a caliper. Then, pictures of equatorial transverse sections of fruits were taken to count the locule number and measure the pericarp thickness, using a Nikon D5300 camera. Image analysis was performed using the ImageJ software (https://imagej.nih.gov/ij/). The number of measurements ranged from n= 50 to n= 200 depending on the number of fruits produced by the different transgenic plants. For leaf surface phenotyping, pictures of full grown leaves were taken using a Nikon D5300 and analysed by intensity threshold filtering. To measure the leaf thickness, images of leaf sections acquired for immuno-labelling experiments were used with three measurement for each picture (n=70 to 100).

### PD index determination

The localization of FW2.2-YFP at PM and PD was observed using confocal imaging performed on a Zeiss LSM 880 confocal laser scanning microscope equipped with fast AiryScan, using a Zeiss C PL APO x63 oil-immersion objective (numerical aperture 1.4). Staining with FM4.64 at a final concentration of 4 µM was used as a control for PM localization (Bolte et al., 2004). For FM4.64 imaging, excitation was performed at 561 nm and fluorescence emission was collected at 630-690 nm. For YFP imaging, excitation was performed at 514 nm and fluorescence emission collected at 520-580 nm. Staining with aniline blue (Biosupplies, Victoria, Australia) was performed by infiltration of a 0.0125% solution; excitation was performed at 405 nm and fluorescence emission collected at 420-480 nm. The calculation of PD index was determined by calculating the fluorescence intensity of FW2.2-YFP at plasmodesmata and at PM as described (Grison et al., 2019). Images were all acquired with the same parameters (zoom, gain, laser intensity, etc.), and YFP and AB channels were acquired sequentially. Ten to twenty images were acquired with a minimum of three biological replicates. Individual images were processed using ImageJ. A minimum of ten regions of interest (ROI) at PD (using AB as a marker) and in the surrounding PM were manually outlined, and the signal intensity was calculated as the mean gray value (sum of gray values of all the pixels in the selected area divided by the ROI surface) for each ROI.

### Immuno-labelling of callose

The level of callose deposition was determined in leaves and in the pericarp of fruits harvested at 5 and 15 DPA. Leaf fragments were fixed with a 4% formaldehyde solution in 1X PBS for 30 min, using vacuum infiltration (∼100 kPa). They were then embedded in 6% SeaKem® LE agarose (Lonza, Basel, Switzerland), and sections of 100 µm were realized using a vibrating blade microtome (Microm 650V; Thermo Fischer Scientific, Walldorf, Germany). Equatorial pericarp fragments were fixed using the same protocol. Pericarp sections of 80 or 150 µm were prepared, and fixed once more in fresh formaldehyde solution for 30 min, rinsed and kept in 1X PBS until use. The leaf and pericarp sections were then processed using the same protocol. The sections were deposited into a small basket containing MTSB buffer (50 mM PIPES, 5 mM EGTA, 5 mM MgSO_4_, pH=7) to perform the immuno-labelling of callose using the InsituPro VSi automated immunohistochemistry device from Intavis (Köln, Germany). Leaf and pericarp sections were rinsed 4 times for 10 min with 700 µL of MTSB. The sections were then incubated for 1 h with 700 µL of a 10% (v/v) DMSO/3% (v/v) IGEPAL^®^ CA-630 (Merck, Darmstadt, Germany) in MTSB. After rinsing, pericarp sections were incubated for 2 h in a 5% (v/v) Normal Donkey serum (NDS; Merck) blocking solution in MTSB, and 4 h with 700 µL of a 1/250 dilution of Anti-callose primary antibody (Biosupplies) in MTSB supplemented with 5% (v/v) NDS. The sections were then washed 6 times with 700 µL of MTSB, and incubated for 2 h with 700 µL of a 1/250 dilution of anti-mouse IgG Alexa Fluor^TM^ 555 secondary antibody (ab150106; Abcam, Cambridge, UK) in MTSB + 5% (v/v) NDS. Sections were rinsed 6 times in MTSB and incubated with 1 µg/mL Calcofluor white (Fluorescent Brightener 28 disodium salt solution, Merck, in MTSB). After rinsing, the sections were mounted on glass slides with citifluor (AF1-25) (EMS Acquisition Corp., PA, USA) and the slides sealed with nail polish.

Identical confocal microscope acquisition parameters were used for all the samples. Because of the highly heterogeneous cellular structure of pericarp and leaf, the total signal intensity of each tissue was quantified, and signal intensity values were measured by integrating the gray value of all the pixels above the same threshold. A minimum of six measurements was performed at least on 5 sections from at least three different fruits or leaves from different plants, and the experiment was repeated twice.

During the callose immuno-labelling experiments, leaf thickness, cell perimeter in leaves or fruits have been manually measured following staining with Calcofluor on pictures acquired from confocal microscopy using ImageJ.

### DANS assays

Before proceeding the DANS assay, 4-week-old tomato plants were pre-treated by spraying water (mock) or 10 mM H_2_O_2_, followed by a 2 h incubation. Then eight droplets (∼1µL) of 1mM CFDA (Merck, Darmstadt, Germany) per leaf sample were loaded on the upper (adaxial) surface. Then, the diffusion of the dye was monitored on the lower (abaxial) surface of the leaf, 5 min after loading CFDA, using an Axiozoom stereomicroscope V16 (Carl Zeiss Microscopy) equipped with a Zeiss Plan-Neofluar 0.5x (NA 0.19) objective lens, a fluorescence lamp (Lumencor Sola LED) and a GFP-BP filter cube. Several leaves with the same size were used from at least 4-5 plants (n=100). Imaging was performed at the same magnification, laser power and gain and pictures were acquired using a CMOS Axiocam 105 color camera. The CF signal intensity was measured on ImageJ by integrating the signal intensity to the pixel surface.

### Co-immunoprecipitation and mass-spectrometry analysis

Total protein extracts from 100 mg of *35S::FW2.2-YFP* fruit pericarp tissue were prepared using the following buffer: 1X PBS, cOmplete Protease Inhibitor Cocktail tablets (Roche, Mannheim, Germany) and 1% Triton X-100. Samples were incubated in the extraction buffer at 4°C for 30 min with agitation, and then centrifuged (16000*g*, 10 min, 4 °C). The supernatant containing the resuspended proteins was used for immunoprecipitation assay using anti-GFP microbeads provided in the μMACS Epitope Tag Protein Isolation Kit according to the manufacturer’s protocol (Miltenyi Biotec, Bergisch Gladbach, Germany). Approximately, 500 μg of soluble proteins were loaded for each co-IP assay.

Fifty µL of the resulting eluate was loaded on a 10% SDS-PAGE acrylamide gel; gel bands were manually cut and transferred to 1.5 mL Eppendorf tubes. Bands were first washed with 500 µl of water and then 500 µl of 25 mM NH_4_HCO_3_. Destaining was performed twice in the presence of 500 µl of 50 % acetonitrile (ACN) in 25 mM NH_4_HCO_3_. Gel bands were dehydrated twice by 500 µl of 100 % ACN, and finally dried at room temperature. Following destaining, proteins were reduced with 500 µl of 10 mM DTT at 56°C for 45 min. The supernatant was then removed and proteins were alkylated with 500 µl of 55 mM iodoacetamide for 30 min. Gel bands were washed twice with 500 µl of 50 % ACN in 25 mM NH_4_HCO_3_, then dehydrated by 500 µl of 100 % CH_3_CN, and finally dried at room temperature. Twenty microliters of a trypsin solution (Sequencing Grade Modified Trypsin, Promega, Madison, USA), at a concentration of 0.0125 µg/µL in 25 mM NH_4_HCO_3_, was added to every gel region and gel bands were kept for 10 min on ice. Fifty microliters of 25 mM NH_4_HCO_3_ were added, and the samples were kept for another 10 min at room temperature. The digestion was performed overnight at 37°C; then peptides were extracted by addition 100 µl of 2% formic acid (FA). Gel bands were extracted twice by addition of 200 µL of 80% ACN and 2% FA. After solvent evaporation in a Speed-vac, peptides were resuspended in 10 µl of 2% FA, then purified with a micro tip C18 (Zip-Tip C18 Millipore Corporation Billerica MA, USA). Peptides were eluted with a solution containing 2% FA (v/v) and 80% ACN (v/v) and dried until total evaporation. Peptides were resuspended in 7 µl 2% FA before LC-MS/MS analysis.

The LC-MS/MS were performed using the Ultimate 3000 RSLC nano system (Thermo Fisher Scientific Inc, Waltham, MA, USA) interfaced online with a nano easy ion source and the Exploris 240 Plus Orbitrap mass spectrometer (Thermo Fisher Scientific Inc, Waltham, MA, USA). The samples were analysed in Data Dependent Acquisition (DDA). The raw files were analysed with MaxQuant version 2.0.3 using default settings. The files were searched against the *Solanum lycopersicum* genome (ITAG4.1_release January 2022 https://solgenomics.net/organism/solanum_lycopersicum/genome 34689 entries) added with the FW2.2-YFP. Identified proteins were filtered according to the following criteria: at least two different trypsin peptides with at least one unique peptide, an E value below 0.01 and a protein E value smaller than 0.01 were required. Using the above criteria, the rate of false peptide sequence assignment and false protein identification were lower than 1%. Proteins were quantified by label-free method with MaxQuant software using unique and razor peptides intensities (Cox et al., 2014). Statistical analyses were carried out using RStudio package software. The protein intensity ratio and statistical tests were applied to identify the significant differences in the protein abundance. Hits were retained if they were quantified in at least four of the five replicates in at least one experiment. Proteins with a significant quantitative ratio (*P* < 0.05 or 0.01 with or without Benjamini correction) were considered as significantly up-regulated and down-regulated respectively.

## Supplemental Data

**Supplemental Figure 1.** Characterization of plants expressing FW2.2 fused to YFP.

**Supplemental Figure 2.** RT-qPCR analysis of FW2.2 expression in tomato plants.

**Supplemental Figure 3.** CRISPR/Cas9-induced mutations producing truncated versions of FW2.2/CNR lacking the PLAC8 domain.

**Supplemental Figure 4.** DANS assays in tomato leaves.

**Supplemental Figure 5.** Pericarp cell perimeter in the different transgenic lines.

**Supplemental Figure 6.** Characterization of Callose Synthase genes in tomato.

**Supplemental Figure 7.** C*a*lS expression level in leaves and fruit from WT, *35S::FW2.2* and *CR-FW2.2* lines.

**Supplemental Figure 8.** Allele swapping complementation assays.

**Supplemental Table 1.** List of primers used for constructs and RT-qPCR analysis.

**Supplemental Table 2.** List of Gateway vectors used for constructs.

**Supplemental Data Set 1.** List of proteins identified as interactors of FW2.2/CNR.

## Acknowledgements

This work was carried out with the financial support of the French Agence Nationale de la Recherche (grant no. ANR-20-CE20-0002), the GPR Bordeaux Plant Sciences in the framework of the IdEX Bordeaux University ‘Investments for the Future’ program, and the French Ministère de l’Enseignement Supérieur et de la Recherche (PhD grant to A. Beauchet). Mass spectrometry experiments were carried out using the facilities of the Montpellier Proteomics Platform (PPM, BioCampus Montpellier, France). The microscopy analyses were performed in the Bordeaux Imaging Center, a service unit of the CNRS-INSERM and Bordeaux University, member of the national infrastructure France BioImaging, and supported by the French National Research Agency (ANR-10-INBS-04). The help of Lysiane Brocard is acknowledged. We express our deepest thanks to Isabelle Atienza, Aurélie Honoré and Valérie Rouyère, for taking care of the plant culture in the greenhouse.

## Author contribution

N.B., F.G., E.B., N.G. and C.C. conceived the project and designed the research. A.B. and N.B. performed the research. V.R. performed the IP-MS-MS proteomics experiments. M.G. helped in the callose immuno-labelling experiments using the InsituPro VSi automate. All authors analyzed and discussed the results. A.B., N.B., N.G. and C.C. wrote the manuscript with input from the other authors.

